# Estimating mutual information and Pearson correlation on neural evoked responses

**DOI:** 10.64898/2026.05.21.727057

**Authors:** Anni Hukari, Silvia Federica Cotroneo, Riitta Salmelin

## Abstract

In neural evoked responses, small variations in the timing or duration of responses can be observed when the same functional response is recorded in different trials, different experimental conditions or by different sensors. These variations limit the ability of correlation-based methods to detect similarities between signals. Mutual information (MI) provides an alternative similarity measure, capable of capturing both linear and non-linear dependencies, yet its practical use is hindered by lack of consensus on estimators for continuous data and the limited understanding of the behavior of the estimators on realistic signals.

In this work, we investigate how to estimate the similarity of neural evoked responses by systematically comparing sample Pearson correlation with three of the most common MI estimators. We describe their behavior using both simulated signals and real magnetoencephalographic data. In the simulations, the estimators are tested against a set of transformations that depict realistic changes in neural evoked responses. Subsequently, we propose guidelines for defining adaptive lower bounds on the similarity estimates and analyzing the similarity rankings induced by the different estimators.

Our findings reveal trade-offs between measures sensitivity and different signal properties. We confirm that Pearson correlation is reliable in describing linear relationships for low-noise signals, and we identify parameter settings that stabilize MI estimators, enabling them to capture complex signal dependencies. Together, these results introduce practical parameter choices and thresholding strategies for mutual information and provide guidance for selecting and interpreting similarity measures in the analysis of neural evoked time series.

## Introduction

When the brain responds to the same stimulus across different trials or experimental conditions, the resulting neural activity is typically considered consistent. However, in this consistent activity, small variations — such as in the timing, duration, or background noise — often occur (Arieli et al. 1996). When analyzing time series in neuroscience, for example those from neurophysiological recordings like electroencephalography (EEG) or magnetoencephalography (MEG), it is useful to define a measure of similarity between signals, able to account for these differences.

Similarities between signal time courses are generally thought to reflect shared behavior across different systems. As multi-subject analysis compares neural responses in different subjects to identify group-level effects, functional connectivity compares signals from different brain areas to find connections between them. On top of these comparisons, neuroimaging signals are also regularly compared with time courses of additional measurements, such as a stimulus time course or another physiological signal, to reveal links between brain activity and specific external factors.

The most commonly used measure to quantify pairwise similarities between time series is Pearson correlation. As a model-based measure of similarity, it is often applied under the assumption that the two input signals are jointly Gaussian and free of significant outliers. When these assumptions hold, Pearson correlation is easy to estimate from experimental recordings — typically as their sample Pearson correlation coefficient — and offers a straightforward interpretation. However, when the assumptions fail, or the signals share more complex dependencies beyond just linear ones, Pearson correlation becomes less reliable and harder to interpret. Mutual information (MI) offers an alternative, model-free measure capable of capturing both linear and higher-order relationships between two input signals (Shannon 1948; Kolmogorov 1956). These features make MI a powerful tool for exploring signal similarity, especially in complex, real-world data. However, unlike Pearson correlation, mutual information has no simple, universally accepted estimator for continuous data, and its accurate estimation from limited samples — common in neurophysiological recordings — is challenging. Several estimators have been developed, e.g., Fraser and Swinney (1986); Darbellay and Vajda (1999); Moon et al. (1995); Kraskov et al. (2004); Cellucci et al. (2005); Ince et al. (2017); Belghazi et al. (2018), each with its advantages, disadvantages, and biases. In all these cases, though, factors such as the underlying probability distribution of the data, the choice of the estimator-specific parameters, and normalization factors can all highly influence their estimates.

Despite these estimation challenges, mutual information has found application in analyzing neuroimaging signals, as they can often be noisy, non-Gaussian, and linked by interesting non-linear relationships. MI is used to analyze continuous signals from neurophysiological recordings, such as electroencephalography (EEG) and magnetoencephalography (MEG), and a common application is the estimation of the similarity between signals recorded from different parts of the brain (Xu et al. 1997; Chen et al. 2000; Jeong et al. 2001; Na et al. 2002; Alonso et al. 2011; Bonita et al. 2014; Yin et al. 2017; Sun et al. 2018; Thuraisingham 2018). In addition to comparing different signals originating inside the brain, mutual information can also be used to detect and remove artifacts — by comparing neural and artifactual signals, e.g., Abbasi et al. (2015, 2016); Hoffmann and Falkenstein (2008); Tuomaala et al. (2025) — and to isolate stimulus-related components by linking neural activity to stimulus parameters (Panzeri et al. 2008; Rousselet et al. 2014; Ince et al. 2015). Additionally, MI is also applied to compare signals across different imaging modalities (Panzeri et al. 2008; Ostwald et al. 2010, 2011a,b). Despite its usefulness, there is no consensus on the best estimator in these contexts, and the existing studies vary widely in the strategies applied to estimate MI, and in general in the way that these values are interpreted. A better understanding on how MI estimators work in different circumstances is still required.

While general comparisons of mutual information estimators have been studied (Papana and Kugiumtzis 2009; Khan et al. 2007; Ross 2014; Czyz et al. 2024), their scope is far from our desired applications. For this reason, our work focuses on the performance of mutual information estimators in in the case of neural evoked responses. Specifically, we evaluated how estimates of Pearson correlation and three widely used MI estimators – namely a binning estimator, the Kraskov–Stögbauer–Grassberger estimator (Kraskov et al. 2004), and a Kernel Density Estimator (Moon et al. 1995) – behave on neural evoked responses, both simulated and originating from a magnetoencephalographic recording. In our simulations, we inspected how the different estimators performed when comparing semi-identical responses, i.e., responses that were identical up to added noise, or a realistic transformation. Among our transformations, we tested different realistic noise conditions, presenting either higher environmental noise, stronger muscle-like noise, or different cropping strategies. We further investigated the effect of differences in the reponse timing, duration or amplitude. Finally, we tested how much the similarity metrics depend on the sampling frequency of the input signals. After exploring these scenarios in the controlled simulated data, we provide an example application on data originating from a magnetoencephalography recording, with the intent to show how the results from the simulations can help us predict the behavior of the estimators applied to real data. Overall, both in the real and simulated data, we focused on how the estimator choice affected the resulting similarity estimates, and whether these outcomes remained interpretable.

It is indeed important to note, when evaluating the performance of similarity estimators, that their reported values cannot be directly compared across different estimators. For this reason, we chose not to focus on comparing their estimated values in absolute terms. Neither Pearson correlation nor mutual information provides an absolute measure of the relationship between two signals. Rather, they serve as tools to establish an order relation — for example, determining whether “Signal A is more related to Signal B than to Signal C.” To use this property, we observed the similarity ranking induced by the estimators rather than the individual reported similarity values. We further leveraged this concept by defining an adaptive minimum-similarity (lower) bound for each estimator and analyzed how different input signals scored relative to this reference case. Although lower bounds in the case of PCC are common practice, this concept is not commonly adopted in the case of mutual information, due to the different nature of the similarity estimate.

In conclusion, this study provides a deeper understanding of how different similarity metrics — and, crucially, their estimators — behave when applied to neurophysiological time series. We introduce a systematic framework for interpreting similarity estimates using adaptive lower bounds. Based on this framework, we offer practical guidance on when each estimator is most reliable, taking into account both signal properties and parameter choices.

## Methods

In all of our analyses, we analyze pairwise similarity between two time series, and treat these two input signals as realizations of the continuous random variables, *X* and *Y*. Both PCC and MI are hereafter defined with respect to this notation. The following subsections explain in detail how the similarity measures can be estimated when *X* and *Y* are finitely sampled, i.e., discretely sampled in time.

### Pearson correlation coefficient (PCC)

The Pearson correlation coefficient (PCC) between two finite-sampled random variables, X and Y, is estimated as the sample correlation coefficient as follows:

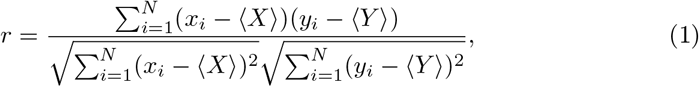

where *N* represents the number of samples, *x*_*i*_, and *y*_*i*_ are the *i*th samples of *X* and *Y*, and ⟨*X*⟩ and ⟨*Y*⟩ represent the mean values, calculated over the samples of the two inputs.

### Normalization

When estimating the Pearson correlation coefficient (PCC), we chose to report only absolute values |*r*|, discarding directionality. For |*r*| = 1, the samples of the two variables are either perfectly linearly correlated or anti-correlated. The opposite case of |*r*| = 0 mean a lack of consistent linear relationship. However, a zero correlation does not imply the absence of structure, as even alternating regions of positive and negative correlation can cancel out.

### Mutual information (MI)

Given two discrete random variables *X* and *Y* with joint probability distribution *p*(*x, y*) and corresponding marginals *p*(*x*) and *p*(*y*), their mutual information is defined as follows:

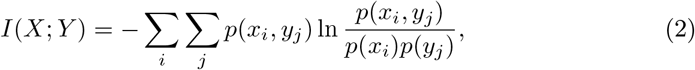

where the sums are taken over all possible outcomes *x*_*i*_ and *y*_*j*_ of the two random variables. Under reasonable conditions (Kolmogorov 1956), this definition can be extended for two continuous random variables, or time series, X and Y as:

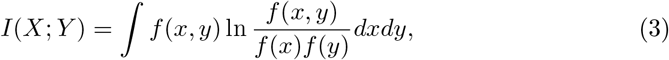

where *f* (*x, y*) is their joint probability density function, and *f* (*x*) and *f* (*y*) are the probability density functions of the two random variables. In this equation the integral is calculated over the range of the two random variables *X* and *Y* and ln(·) is the natural logarithm, yielding values of mutual information in natural units of information (nat). In the case of all the estimators reported here, we express all MI estimates in nats (logarithmic base *e*) for comparability.

To estimate mutual information directly from this definition, it is essential to know the probability density distribution functions of the random variables. These distributions can be very difficult to estimate from real data, due to their finite sampling, and different methods have been proposed to generate estimates for mutual information in this case. The following sections describe the three estimators that were chosen for our analysis and their corresponding parameter choices.

### Kernel Density estimator (KDE)

The kernel density estimator (KDE) is a non-parametric method for estimating the probability density function (pdf) of a sample dataset. Estimation relies on fitting a kernel function with a fixed width — or bandwidth — on each data point and estimating the pdf of the dataset by summing all these kernel functions (Moon et al. 1995; Silverman 1998). The method is sensitive both to the choice of kernel function and to its bandwidth. In our analysis, we chose to use a Gaussian kernel as suggested by Moon et al. (1995).

For a Gaussian kernel, the choice of the kernel bandwidth heavily impacts the outcome of the pdf estimate. A smaller bandwidth, relative to the input spread, yields more detailed estimates, but if it is too small, the estimate becomes overly sensitive to local structures and noise. Analytical results, such as Scott’s rule (Scott and Terrell 1987), can be used to guide the choice of optimal bandwidth.

If the kernel bandwidth is defined as the covariance matrix of the input data scaled by *f* ^2^, Scott’s rule gives the optimal value of *f* as:

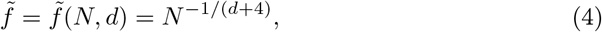

where *N* is the number of samples in the input, and *d* is the dimensionality of the data.

Notably, Scott’s rule defines the optimal kernel bandwidth separately in the case of the estimate of the univariate and bivariate pdfs. Compared to using the same scaling factor for all the estimates, this approach adapts better to the data.

We implemented the KDE estimation of the pdfs using the sciPy library (v1.14.0) (Virtanen et al. 2020) and controlled the bandwidth through the library-defined band-width factor *f*. We kept the bandwidth ratio between the univariate and bivariate distributions the same as in Scott’s rule.

After estimating the probability density functions 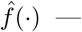both for the univariate and bivariate distributions — with KDE, we further applied the following simplification of Eq. (3) from Steuer et al. (2002) to estimate mutual information:

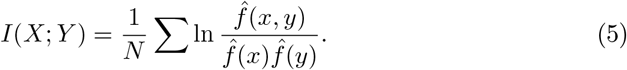

### Kraskov–Stögbauer–Grassberger (KSG) estimator

The Kraskow–Stögbauer–Grassberger (KSG) estimator estimates mutual information directly from the data samples, relying on a nearest neighbor algorithm (Kraskov et al. 2004). The KSG estimator employs the following relationship to estimate mutual information:

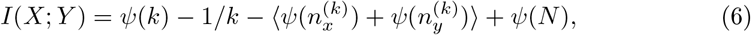

where *ψ*(·) is the digamma function, *k* is the number of neighbors considered, *N* is the number of samples, and 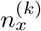 and 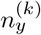 are a derivation — in the univariate domain of *X* and *Y* — of the *k*-nearest neighbors in the bivariate space of (*X, Y*).

The KSG estimator requires the user to define a single parameter, the number of neighbors, *k*, to estimate the mutual information between two random variables. The number of nearest neighbors chosen depends on the amount of precision one expects in the data, but the estimate is reasonably stable with respect to changes in the choice of *k* (Papana and Kugiumtzis 2009; Ross 2014).

We utilized the ennemi Python library (v1.4.0) developed by Laarne et al. (2021) to estimate mutual information with KSG. Prior the estimation, the input variables were scaled to unit variance, as recommended by Kraskov et al. (2004).

### Adaptive Binning (AB)

A different approach to estimating probability density function of a sample is to discretize the data into frequency histograms. Defining the bins by equal occupancy has been shown to work best for unevenly distributed data, such as neurophysiological time series (Ince et al. 2017). This strategy is generally known as adaptive binning (AB). In our implementation, we used this binning strategy on the marginal distributions of the two signals and then applied the binning scheme derived from the marginals to estimate the joint distribution. Mutual information was then calculated from the estimated discretized probability distribution by plugging in the estimates into Eq. (3).

In this binning strategy, the most important parameter to tune is the number of bins *K* used to approximate the distributions. The choice of *K* is dependent on the sample size and can highly impact the estimator results. Analytical results such as Doane’s rule (Doane 1976) can be used to determine this parameter optimally. Doane’s rule states that, given a random variable *Z*, with range *Ƶ*, sampled by *N* samples yielding the expected value ⟨*Z*⟩, the optimal number of bins *K* is defined as:

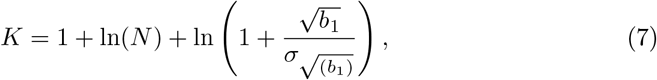

where:

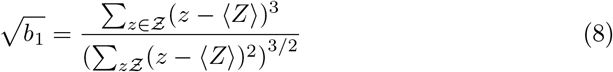

and

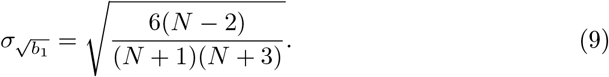

Even with an optimal choice of *K*, binning methods are known to yield upward-biased estimates for mutual information due to the limited sample size *N*. As suggested by Cohen (2014), we estimated and corrected the inflation of mutual information as Steuer et al. (2002):

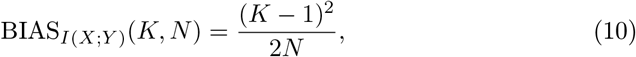

where *K* is the number of bins defined in the univariate distributions, and *N* the number of samples per random variable.

### Normalization

Mutual information is an unbounded measure, with values ranging from 0 to +∞. This broad range of values poses challenges in interpreting and comparing its estimates. To restrict the variability range of our estimates, we normalized the mutual information values, as done, for instance, by Linfoot (1957), as follows:

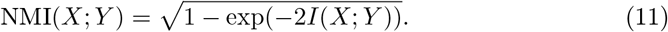

With this normalization, the maximum value NMI = 1 indicates that *X* is perfectly defined by *Y*. This is true in any case where the two random variables are linked by a bijective transformation, including linear correlation (or anticorrelation). Conversely, NMI = 0 implies that the two random variables are statistically independent: nothing can be inferred about one variable from the knowledge of the other one. This choice of normalization depends on the units used to express mutual information and leads to uneven spacing of the normalized values, introducing an upward bias in the lowest mutual information estimates.

### Adaptive lower bounds

To assess the significance of the similarity estimates, we defined adaptive lower bounds for each comparison. Because most analyses kept one variable *X* fixed while parametrically modifying the second variable *Y*, the lower bound was defined relative to *X*. Specifically, we constructed an empirical null distribution by assuming statistical independence between *X* and *Y*. This null condition was approximated by replacing *Y* with Gaussian noise matched to *X* in mean and standard deviation. Similarity estimates were computed for 1000 independent noise realizations, yielding a distribution that represents the expected similarity between *X* and an unrelated signal. Statistical significance was assessed by deriving empirical 99% confidence thresholds (corresponding to *p* = 0.01) from this null distribution. For Pearson correlation, this procedure reduces to testing the standard null hypothesis of zero correlation between the two time series.

### Simulated neural evoked responses

All the analyses presented here were performed using Python, and the code has been made publicly available.

### Simulations

Our simulations included the following steps: 1) a base signal was simulated, and two copies were stored; 2) a transformation was applied to one or both of the two copies; 3) both signals were scaled, and noise was added to each one independently.

For each choice of base signal and transformation parameters, this procedure was repeated with different noise iterations 1000 times. The upcoming sections expand each step of this procedure in more detail.

#### 1. Base signals

The event-related simulated base signals *X*(*t*) were defined as Morlet wavelets such that their shape was controlled. The shape was kept identical across responses by controlling the simulations parameters. More details on the implementation of the simulated Morlet wavelets and the parameters are presented in Simulating evoked neural responses. Unless otherwise specified, the base signals had a characteristic frequency of about 8.3 Hz and were simulated in a time window of 0.35 s and sampled at 600 Hz.

#### 2. Transformations

After simulating one base signal, two copies were stored: *X*(*t*) and *Y* (*t*). The relationship between them was then altered by applying a selected transformation. A summary of these transformations is provided in Table 1.

**Table 1.** Summary of the transformations applied to the simulated event-related base signals.

| Transformation | Description | Real-life analogue |
| --- | --- | --- |
| Sample size (A) | The sampling frequency ($f_s$) was changed identically for both signals. | Variation in the number of measured samples in signals due to different measuring. |
| Noise amplitude (B) | The SNR was changed in both signals by modifying the noise amplitude identically for the two signals. | Contribution of low-amplitude noise in the signals — e.g., sensor or environmental noise — being more or less prominent. |
| Outliers (C) | Outlier point — i.e., points with z-score in a specific range, were added to one of the two signals. | Presence of sparse high-amplitude noise, such as muscular activity, in the responses. |
| Noisy segments (D) | Zero-padding was added at the end of both signals identically. | Variation in the epoching strategy or the length of the measured interval. |
| Timing (E) | $Y(t)$ was obtained by shifting $X(t)$ in time by $\Delta t$: $Y(t) = X(t + \Delta t)$. The shifting ranged from 0 to the case of the two time series no longer overlapped. | Timing differences between the recorded responses due to non-synchrony of the sensors or inherent timing differences in the responses. |
| Duration (F) | The duration $T$ of $Y(t)$ was modified, while both signal peaks were kept at $\bar{t} = 0.1$. | Different durations of the two responses. |

#### 3. Scaling and noise

After applying the transformations, the signals were scaled to constrain their values within the range [−1, 1], while preserving their relative differences. The scaling was achieved by dividing both signals by their shared maximum absolute value. Scaling was applied before adding noise to the signals and, in the case of the outlier transformation, the scaling was done before adding the outlier points.

Random Gaussian noise with zero mean and standard deviation *σ* was added to the scaled responses in all simulations. The standard deviation *σ* was chosen such that the signal-to-noise ratio (SNR) of the noisy signal was within 0.5 dB of a target SNR. In the case of outlier insertion and noise-only segments transformations, the target SNR was calculated with respect to the base signal — before the transformation was applied — to keep the noise levels consistent with the other examples.

## Results

### Parameter choices

As noted in Section 1, each of the three MI estimators depends on the choice of one free parameter, and their estimates are sensitive to how these parameters are chosen. Before the simulation study, we explored broader parameter ranges for each estimator and selected the values used in our simulations.

Testing was conducted under two conditions (Fig. 1). In the first condition (Fig. 1a), two simulated responses differing only by added Gaussian noise (SNR = 20 dB) were compared. In the second condition (Fig. 1b), the reference response was compared with a nonlinearly transformed (quadratic) version of the same signal. For each estimator and condition, lower bounds were computed using identical parameter settings to those used for the corresponding similarity estimates.

**Fig. 1.**
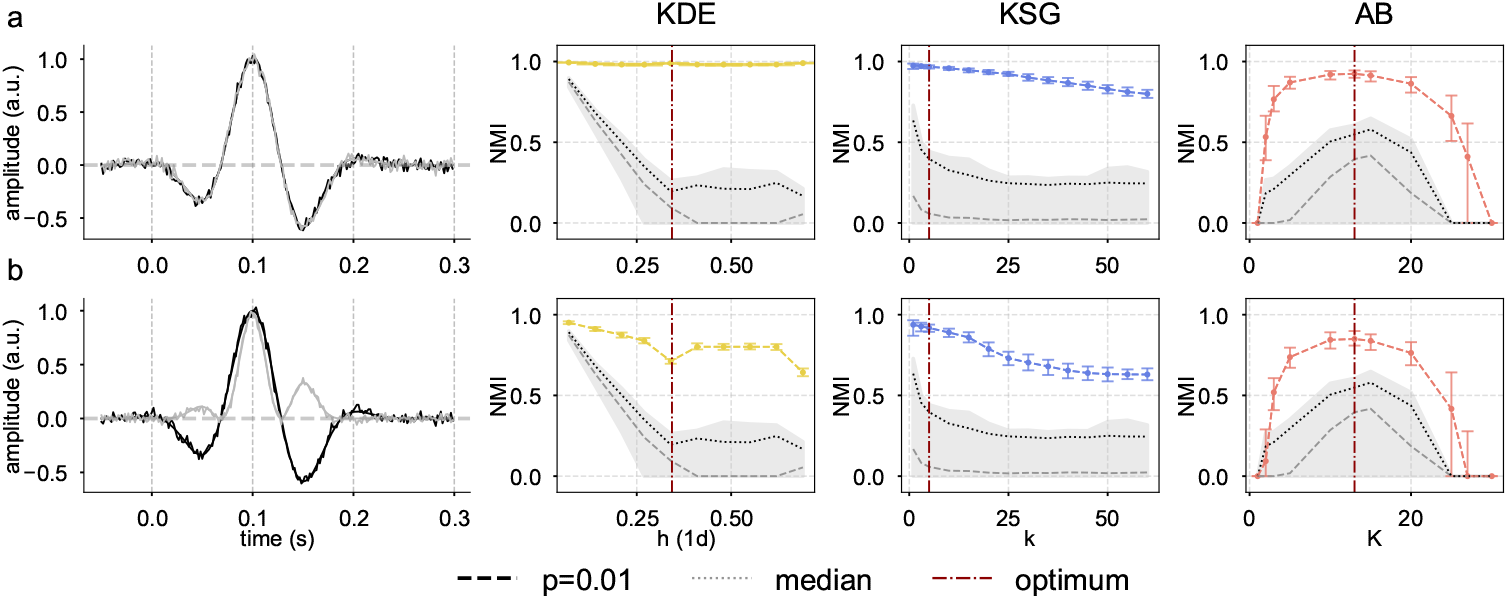
Results of the parameter testing for the three MI estimators in two input-signal scenarios. Column 1 depicts examples of the input signals and columns (2–4) the results of the MI estimators. For each choice of the parameter (x-axis) the comparison was run 1000 times (runs) over different noise iterations. The colored dots represent the median value of the iterations and the bars around it the full variability range of the estimates. The gray shaded area shows the full variability range of the lower bound, with the lighter dashed line indicating the median value and the darker line the edge for the confidence interval of *p* = 99%. The vertical red dashed lines show the final choice for the free parameter in each case. a: Comparison of two semi-identical input signals, differing only by the noise iteration, which varies for each run. b: Comparison of two signals related by a quadratic relationship. Each run has a different noise iteration.

Overall, we found that parameter choices systematically affected the estimates, often biasing them toward lower values. This effect was evident when the two inputs were nearly identical (first scenario) and became more pronounced when the relationship was strong but nonlinear (second scenario). The lower bounds proved useful for identifying the optimal parameter choice.

KDE was relatively insensitive to bandwidth choice when comparing semi-identical signals, but became more sensitive in the more complex comparison scenario. The lower bounds made this effect clearer: small bandwidths produced high lower-bound estimates, making it difficult to distinguish true signal similarity from noise. Increasing the bandwidth reduced the lower bounds and improved separability between signal and noise comparisons. Scott’s rule yielded stable performance in both respects, producing high similarity estimates for related signals while maintaining a clearly separated lower bound, and was therefore selected for all subsequent analyses.

For KSG, the estimates of similarity between the inputs decreased as the parameter *k* increased in both scenarios. For very small values of *k*, the estimates were highest, but the corresponding lower bounds were also elevated, reducing separability between signal and noise comparisons. Starting at *k* = 5, the gap between the two comparisons widened, while keeping a high similarity estimate between the two statistically dependent inputs. Although Kraskov et al. (2004) recommend choosing *k* between 2 and 4, we selected *k* = 5 to improve the distinction between signal and noise comparisons.

For AB, the effect of varying the number of bins was more complex. Unlike the other estimators, both scenarios showed similar trends, and the optimal value of *K* corresponded to the peak of the similarity estimate curve. At this value, the estimates were closest to the expected similarity in both scenarios, while the lower bound remained sufficiently low to preserve separation from noise. Doane’s rule correctly identified this optimal bin number (13 in our case) and was therefore adopted in our analyses.

### Simulations

After fixing the parameters for all the estimators, we investigated the impact of different signal properties on the similarity estimates through simulated parametric signal manipulations. The main results of our analyses for the four estimators are presented in Fig. 2.

**Fig. 2.**
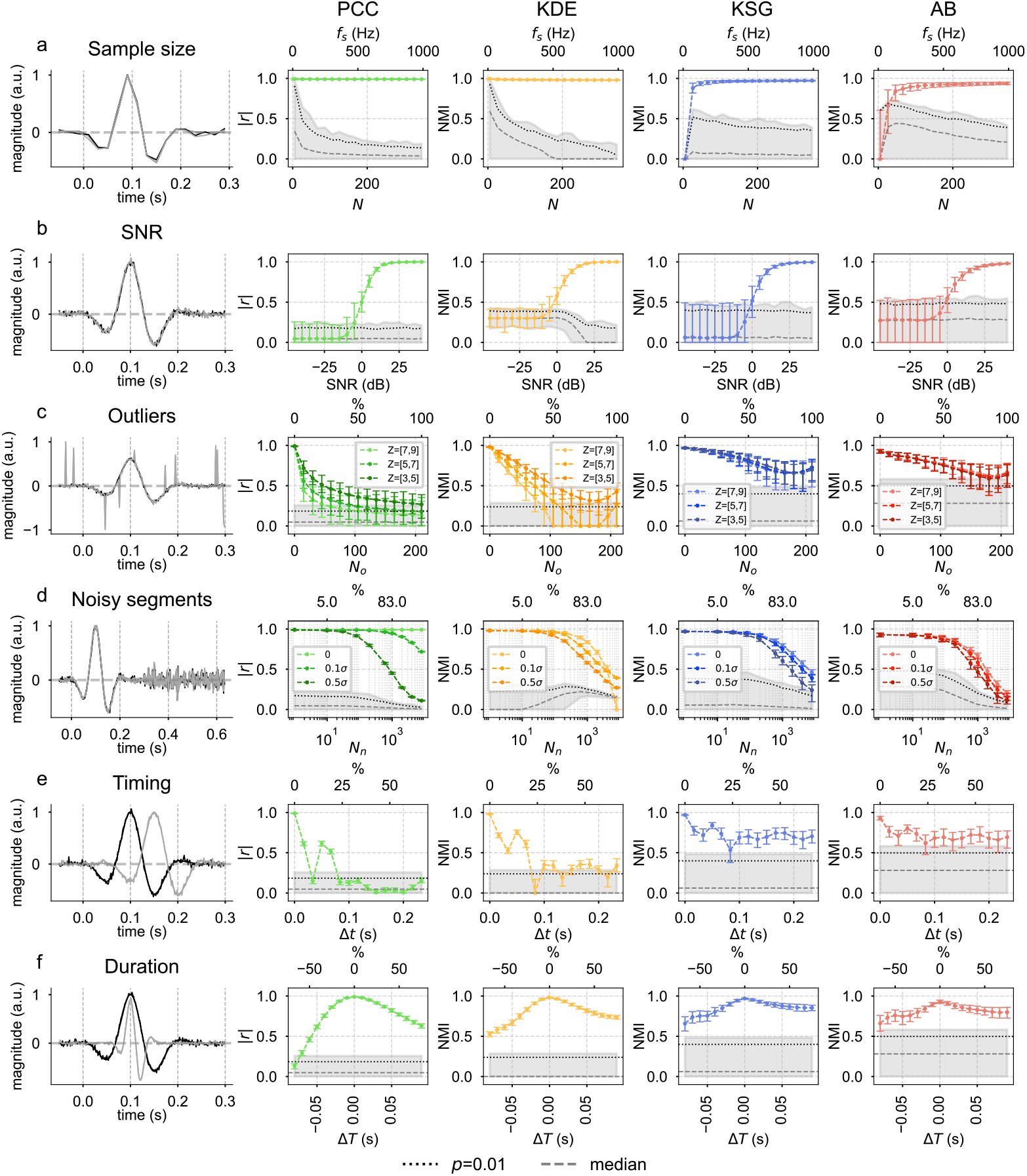
Similarity estimates on simulated data. Column 1 reports a visual example of the input signals after the given transformation (as described in Table 1); each transformation is controlled by a free parameter. Columns (2–5) show the results of the different similarity estimators over the parametric analysis of the transformed signals. For each choice of the parameter (x-axis) the comparison was run 1000 times over different iterations of noise of the input signals. The colored dots represent the median value of the iterations and the bars around it the full variability range of the estimates. The shaded area shows the full variability range of the lower bound, the lighter dashed line represents the median value and the darker line the edge for the confidence interval of *p* = 99%.

### Sample size

The first parameter examined in the simulations was the robustness of the similarity estimators to input size (Fig. 2a). As expected, all estimators returned high similarity values when comparing semi-identical signals once sufficient samples were available. For PCC and KDE, similarity values remained high even at very small sample sizes (*N* < 6), but were indistinguishable from those obtained when comparing signals with pure noise. In contrast, AB and KSG produced lower similarity estimates at small sample sizes, but again, at very low sample sizes their values largely overlapped with the corresponding lower bounds.

For all four estimators, the gap between semi-identical and noise comparisons widened as sample size increased, showing that larger sample size improves the reliability of the similarity comparison. At the same time, excessively large sample sizes were more likely to create near-duplicate samples, a known negative factor for the KSG and KDE estimators (Laarne et al. 2022; Moon et al. 1995). Overall, even at relatively low sample sizes (≈100), all estimators yielded values which were distinct from their lower bounds and eventually converged to estimates with similarly low spreads with at least 210 samples, corresponding to a sampling frequency of 600 Hz, which was then used in the following examples.

### SNR

Next, we examined the effect of noise amplitude (Fig. 2b). All estimators struggled to separate semi-identical signals from noise when the inputs’ SNR was below 0 dB, but performance improved as the SNR increased. PCC performed slightly better, separating the comparison of the two signals from the one of the base signals vs. noise, already at SNR = 0 dB. The MI estimators, on the other hand, required positive SNR values. In all cases, median values rose, and variability decreased with increasing SNR, stabilizing around 20 dB, which we adopted for our subsequent simulations. Among the estimators, KSG and AB showed a more pronounced spread at low SNR compared to PCC and KDE. KDE notably failed at very high SNR (> 150 dB) due to the presence of singularities in the covariance matrix when nearly identical samples occurred. Both KSG and KDE are known to require some noise in the inputs to be defined properly: for KSG, it helps to mitigate autocorrelation (Laarne et al. 2022; Moon et al. 1995), whereas for KDE it prevents degeneracy of the kernel covariance.

### High-amplitude sparse noise

We then tested the effects of outliers (or high-amplitude sparse noise) by adding them to one of the two semi-identical responses and varying their amplitude across three *z*-score ranges (Fig. 2c). In this case, PCC and KDE were the most affected: their median values dropped quickly as outlier amplitude increased, often falling below their respective lower-bound threshold. KSG and AB were more robust, showing a slower decline. Across all estimators, the spread of values widened as the proportion of outliers grew, a trend especially pronounced for the MI-based methods. Interestingly, all MI estimators showed a slight rebound in median values when about 80% of the points in one signal were outliers.

### Signal cropping

Because neural evoked responses contain segments with varying levels of activity, cropping affects both the amount of retained information and the effective signal length. To model this, we compared two semi-identical responses and appended noise-only segments with different variances, expressed relative to the standard deviation of the original signal (Fig. 2d). Except for PCC in the noiseless case, all estimators showed decreasing similarity values as the proportion of noise-only points increased. This effect became most pronounced when non-informative segments exceeded 50% of the time series, with steeper declines and larger variability for MI estimators than for PCC. Although PCC values approached the lower bound as noisy segments increased, they never dropped below it. In contrast, MI estimators exhibited simultaneous decreases in median values and increases in the spread of their estimates. MI estimates reached the lower bound only when nearly the entire signal was replaced by high-variance noise. Importantly, for all estimators the lower bounds decreased with increasing signal length, indicating that smaller similarity values can remain statistically significant for longer signals.

### Time shifts

For time shifts (Fig. 2e), PCC and KDE values fluctuated strongly and dropped below the lower bound even for small shifts. In contrast, the KSG and AB estimators were more robust, typically remaining above their lower bounds, especially for modest shifts. While median values remained high, the spread of the estimates was larger than for PCC, especially in the case of KSG and AB. The dependence on shift magnitude was non-linear, with three local maxima corresponding to alignments of positive or negative peaks. Overall, AB and KSG exhibited similar patterns and generally remained above the lower bound, whereas KDE behaved more similarly to PCC but stayed above the lower bound until the two inputs showed minimal temporal overlap.

### Duration

When the signal peak timing was fixed but the duration was altered, all estimators returned values above their lower bounds but showed asymmetry. Shortening a signal reduced similarity more than lengthening it, since shrinkage introduced more noise-only segments. This effect was strongest for PCC, whose median values fell close to zero, dropping below the lower bound for large shrinkage. The three MI estimators displayed a milder decline, though AB and KSG showed higher variability than KDE, especially as duration differences increased.

### Magnitude

On top of these transformations, we tested if we could find any contribution given by changes in the relative magnitude of the two semi-identical signals but none was found in the different estimators, except in the case of KSG. As mentioned in its imprementation, this estimator requires standardization of the input signals, as stated in its implementation (Laarne et al. 2022).

### Case study using MEG data

To test how the patterns observed in the simulations generalize to real data, we applied the four similarity estimators to MEG recordings from a picture-naming task (Ala-Salomäki et al. 2021). Data were preprocessed as in the original study, epoched around stimulus onset, baseline-corrected. The resulting epochs were averaged in each channel to obtain estimates of the evoked responses. Following the simulation results, we fixed the signal length at 510 samples (sampling frequency 600 Hz), added minimal Gaussian noise so that the KDE and KSG estimators would stabilize, keeping an SNR of 20 dB.

Whole-head MEG records magnetic activity using an array of channels arranged as a helmet around the head. An individual magnetic source can contribute to the signals picked up by several channels, due the geometry of the generated magnetic fields. For this reason, different channels can record highly dependent signals, not only capturing common neural activity, but also shared noise which can stay in the data even after preprocessing it.

To illustrate the estimators’ behavior on realistic data, we present the result from a single subject from the original study. We compare one reference channel — chosen near a known brain activation source — with all other sensors (Fig. 3a–b). Comparable patterns were observed in other subjects.

**Fig. 3.**
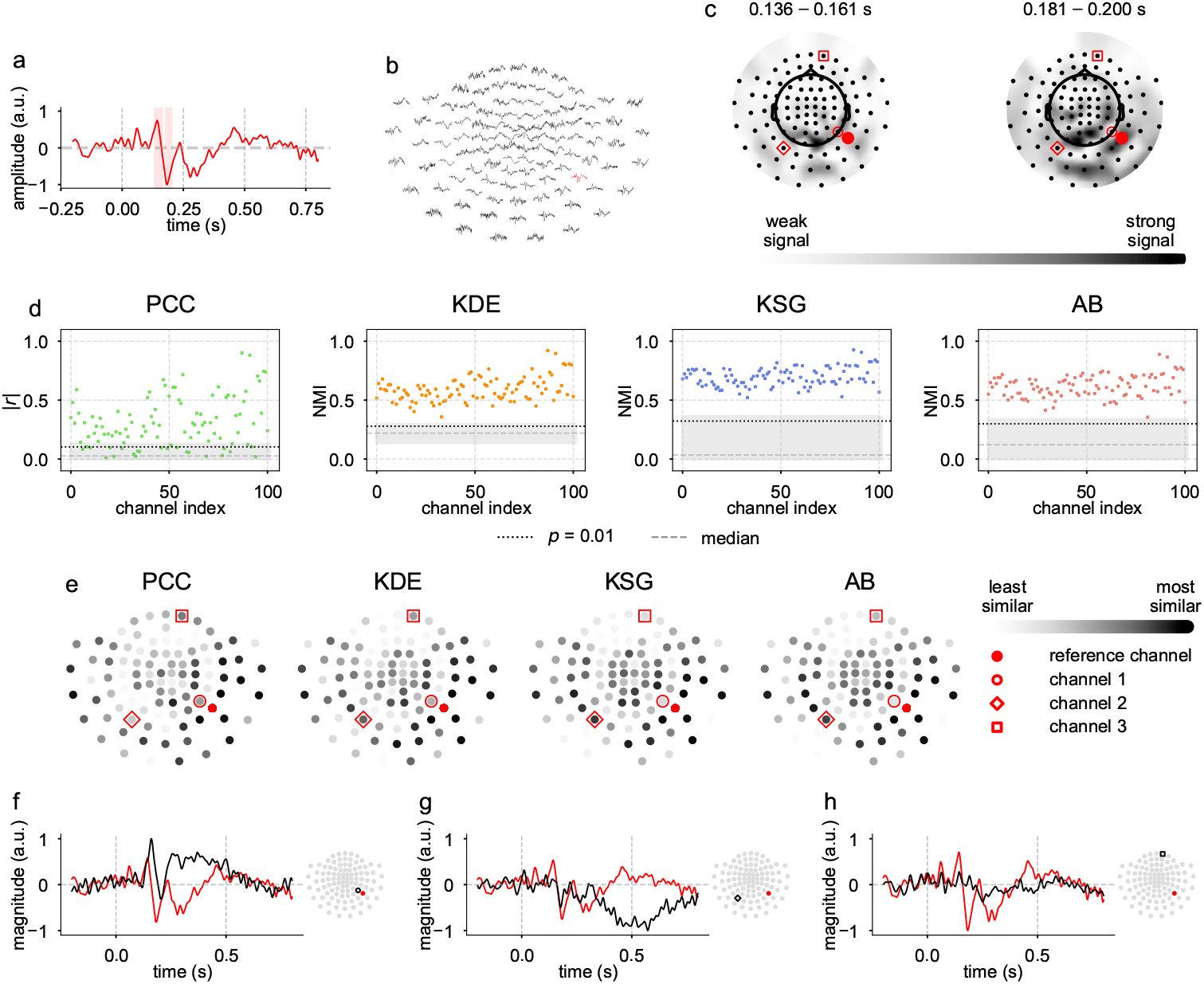
Results of the similarity estimators on recorded magnetoencephalographic data. a: The preprocessed and standardized signal from the reference channel. Shaded areas represent time windows of interesting activation. b: The recorded MEG signals displayed on a 2D projection of the MEG channels. The channels shown are planar gradiometers (longitudinal orientation) from a MEGIN Triux neo MEG device. The signal reference channel is highlighted in red; those from the other channels are shown in black. c: The spatial maps of the signal RMS, averaged at the given time windows. The tme windows were chosen around the main peaks in the reference channel signal (shaded areas in Panel a). The RMS of the channel signals are averaged in the given time window, interpolated on the surface of the MEG helmet and displayed on its 2D projection. Black dots mark channel locations, and red contour identifies the channels of interest investigated in Panels f-g; black/gray shading indicates the interpolated RMS of the channel signals. d: The similarity estimates between the reference channel and all other channels. Results for each estimator are in the different columns. The colored points show the pairwise comparisons; the gray shaded area indicates the lower bound, with median (dashed lighter gray) and 99% confidence limit (dashed dark gray). e: The similarity rank maps. Shading indicates normalized channel similarity with respect to the reference (red). For each estimator, values are normalized between its minimum and maximum. f-–h: Example of pairwise comparisons. Panel f shows a case where PCC shows higher similarity ranking than the MI estimators for a channel close to the reference; Panel g shows another case where the MI estimators rank the similarity between the two channels higher than PCC; panel h shows a case where all estimators rank high similarity with a frontal distant channel, possibly sharing an artefactual component with the reference channel.

We computed the similarity estimates between the reference signal and signals from all other channels (Fig. 3d). Analogue to the simulation study, we defined lower bounds by comparing the reference signal with 1,000 Gaussian noise realizations matched in standard deviation. The four estimators were again split in two behavioral pairs. PCC and KDE on one hand yielded more variable estimates and tighter lower bounds, while KSG and AB, on the other, produced broader bounds and more uniform similarities across sensors. For PCC, several comparisons fell below its lower bound, indicating no detectable linear relationship higher that with pure noise. In contrast, all MI estimators produced similarity values significantly above their respective lower bounds, confirming that all the MEG signals from the same recording share significant information. This information is due to their common stimulus-driven origin and the fact that all channels have been recorded under the same noise conditions and underwent the same preprocessing.

Since all signals shared so much information, only referring to the comparison with random noise provided limited insights. However, the magnitude of these estimates still varied for the different pairwise comparisons. To better understand the relative importance of the channels, we ranked them with respect to their similarity estimates (Fig. 3e). Across all estimators, sensors closest to the reference showed the highest similarity ranks, consistent with the spatial spread of the underlying magnetic sources. Beyond this shared pattern, differences emerged: PCC and KDE tended to rank distant but linearly correlated sensors higher, whereas KSG and AB assigned higher similarity also to sensors capturing delayed or slightly distorted responses. Some examples of this behavior can be seen in the activation maps (Fig. 3c), where channels ranked as highly similar by KSG and AB are shown to be measuring a stronger signal at the same time window as the reference channel. These examples confirm how PCC favors strict linear correspondence, while MI estimators can detect nonlinear, or piecewise alternating linear and antilinear activity (Fig. 3g). Some examples of the discrepancies between estimators, or similarities which could not be explained by just looking at the significant signal intensities, are presented in Fig. 3f–h.

Using a reference signal recorded under the same conditions as the other channels may partly limit the neuroscientific interpretability of the results, since shared artifacts (e.g., eye blinks) can inflate similarity values in channels which may not capture interesting brain activity (e.g., frontal ones) even after preprocessing. Nevertheless, the general ranking patterns and estimator behavior remain interpretable.

## Discussion

Starting from three mutual information estimators and the sample estimator for Pearson correlation, this study examined how parameter settings and signal properties affect their suitability for analyzing neural evoked responses, or other similar electrophysiological data. We emphasize the usefulness of ranking similarity values rather than interpreting absolute magnitudes and propose using adaptive lower bounds as references, also in the case of real data.

As shown in previous work, all MI estimators are sensitive to biases due to the input signals, such as high autocorrelation (Steuer et al. 2002), hard-to-estimate probability distributions (Kraskov et al. 2004), presence of identical samples (Kraskov et al. 2004) and limited number of samples (Gao et al. 2015). On top of these, mutual information normalization also adds further biases, and the normalization strategy presented in our analyses has been reported to introduce an upward bias (Nagel et al. 2024). Our simulations confirmed all of these biases and limitations but also pointed to practical strategies for stable performance across realistic signal shapes. More generally, all estimators required appropriate parameter choices, sufficient sample size, and adequate signal-to-noise ratio. Without these, results became unstable and not distinguishable from noise.

For the signals shapes presented in this study, the minimum number of input samples differed for the different estimators. AB was found to be the most sensitive to small input sample sizes making it the least suitable in cases when a limited number of samples is available. In general, for KDE and PCC, smaller sample sizes lead to overestimation of the similarity, and for AB and KSG, instead, to underestimation. In all cases, too small sample sizes made the comparison of two highly similar signals indistinguishable from the comparison with pure noise. For this reason, we believe that defining such lower bounds is especially important when interpreting similarity estimates. Overall, sampling frequencies over 200 Hz, corresponded to a sufficient signal coverage and reasonable results for all the similarity estimators for the signals presented in this study. Increasing sampling frequencies up to 500 Hz improved the stability of the estimates, but oversampling at even higher frequencies did not bring significant improvements in the results. Too high sample sizes, instead, increased computational cost and the probability of introducing biases due to sample-redundancy and autocorrelation. These results were confirmed both in our simulation study and in our real data example. Good sampling frequencies are easily achievable in common neuroimaging setups, making them a reasonable choice for real data applications. Optimal sampling frequency cannot be defined a priori, as the length of the input signals impacts on the required sampling frequencies: higher sampling frequencies are required with shorter signals to provide enough samples for the estimators to function correctly.

The impact of noise was also characterized, and we found how the stability of all estimators improved as the signal-to-noise ratio of the input signals increased, but the effect plateaued around 20 dB in both simulated and real data. Low levels of added noise were sufficient to stabilize KDE and KSG, and, by keeping the SNR around 20 dB, the added noise caused negligible distortion. Based on these findings, we recommend adding a small amount of Gaussian noise to the input signals — even after preprocessing — when computing similarity estimates with KSG or KDE.

After stabilizing the estimates, generalizing thresholding of the similarity estimates can be a difficult task. While PCC is often thresholded, no consensus exists for mutual information. As we found that not only normalization, but also intrinsic signal properties such as sample size strongly affected similarity estimates, absolute cut-offs become an unreliable threshold. We propose defining adaptive lower bounds for the similarity estimates, by choosing comparisons for which no similarity should exist, but the relevant signal properties — such as sample size and signal variance — are retained. On top of this, we claim that ranking, rather than absolute thresholding, can provide a more interpretable way to read similarity results.

By setting the comparison against adaptive lower bounds, we found MI estimators, especially AB and KSG, to be more prone than PCC to pinpointing relationships in signals subject to certain common transformations. These transformations included: small relative time-shifts in the inputs, small changes in the signal duration and the presence of sparse high-amplitude noise (outliers). The MEG case study validated the behaviors outlined in the simulation study in the case of real data. Importantly, unlike PCC, MI estimators, in general, always indicated that signals belonging to the same recording, as responses to the same stimulus, were significantly more similar to one other than to noise.

Evolving from the comparison with only the lower bound, in the MEG case study, we showcased how important it is to look at similarity ranking rather than individual values. Evaluating how the different estimators ranked the signals similarities highlighted more characteristic behaviors of the estimators.

When analyzing the similarity rankings of all the channel data, with respect to a chosen reference channel, channels ranking the highest for KSG and AB aligned well with experimental activation patterns at times also slightly different from the main activation peak of the reference channel. PCC and KDE also highlighted logical patterns in their similarity ranking but they tended to mainly outline the shape similarity to the reference channel rather than detecting other stimulus-related responses. This behavior caused their overall rankings to differ, partially, from that of KSG and AB, reinforcing the previous result that the two sets of similarity measures are more sensitive to different patterns in the data.

Our real-data example underlined the importance of choosing a good reference signal for the desired application. In our case, we chose a reference channel which contained a mix of neural activity, noise, and artifacts. For this reason, the similarity estimators — especially the MI ones — recognized similarities in the other channels based on all these factors. This behavior can make it difficult to interpret whether high similarity across signals reflects true neural activation or just shared noise or common residual artifacts.

## Conclusions

To sum up the overall behaviors of the estimators, both in the simulations and in the analysis of real data, we found that the four estimators clustered into two pairs with comparable behavior: PCC with KDE, and KSG with AB. Our implementation of the KDE estimator showed some differences from the results of PCC but did not add much to its results and was harder to tune than the other MI estimators. KSG and AB diverged more from PCC, but this different behavior also included a higher variance in their estimates, and more pronounced upward biases especially for the lower values. Between these two, KSG was less sensitive to the parameter choice and the number of available samples, making it the more practical choice between the two. In general, when the comparison aimed at recognizing the exact same shape in different signals, PCC and KDE were found to be more efficient, with PCC being the simpler solution. On the other hand, KSG and AB, more prone to assigning high similarity to signals sharing information in a broader sense, are more suitable for cases in which we do not expect the comparison signals to be identical but still share many features. Among the two, we believe that KSG proved the simpler to tune.

In conclusion, when interpreted correctly, mutual information offers a versatile and powerful alternative to Pearson correlation in analyzing neurophysiological time series, and we believe that the case study presented here can be used as inspiration for future applications. When properly implemented, the estimators presented in this study are reliable tools for quantifying signal similarities, making them valuable for a range of neuroscience applications, and especially in the study of EEG and MEG event-related signals.

## Acknowledgements

We would like to thank Mia Liljeström and Wouter Marijn van Vliet for the helpful comments and discussions regarding our work, and Oliver Kostamo for the interest and contributions in the technical details of our estimator’s choice.

## Declarations

### Funding

This research was funded by Research Council of Finland (grant #355407 to R.S.), the Sigrid Jusélius Foundation (to R.S.) and Jenny and Antti Wihuri Foundation (to S.F.C.).

### Competing interests

I declare that the authors have no competing interests as defined by Springer or other interests that might be perceived to influence the results and/or discussion reported in this paper.

### Ethics approval and consent to participate

The original authors collected the data after obtaining a written informed consent from all participants, in agreement with the prior approval of the Aalto University Research Ethics Committee.

### Consent for publication

All authors have approved the manuscript and agree with its submission.

### Data availability

In agreement with the ethical permission and national privacy regulations at the time of the study, the raw MEG data cannot be made openly available.

### Materials availability

All of the material is owned by the authors and no permissions are required.

### Code availability

The code used to generate the simulated data is publicly available at https://github.com/AaltoImagingLanguage/micorr.

### Author contribution

S.F.C. and A.H. contributed to conceptualisation, methodology, software, validation, formal analysis and writing (original draft + review and editing). A.H. was responsible for visualisation. R.S. contributed with supervision and writing (review and editing). All authors approved the final manuscript.

## Simulating evoked neural responses

The event-related simulated signals *X*(*t*) were defined as Morlet wavelets as follows:

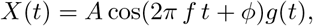

where *A* is the amplitude of the signal, *f* its frequency, *ϕ* its phase, *t* is the time parameter expressed in seconds, and the envelope function *g*(*t*) is:

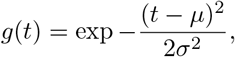

where *µ* is the center and *σ* the standard deviation. The constraints used to setfix the shape of the signals were: the relationship between the center *σ* and the frequency *f* was defined as *σ* = *a/f*, where *a* = 0.36, and the phase shift 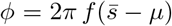, where 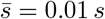. The timing of the main response peak was at the fixed time 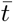 which, unless otherwise specified, was set to be 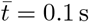. The frequency of the signal was determined by the period, or duration *T* = 1*/f*, which, unless otherwise specified, was set to be *T* = 0.12 s.

## Notes

https://github.com/AaltoImagingLanguage/micorr

### Competing Interest Statement

The authors have declared no competing interest.

## References

Abbasi, O., Dammers, J., Arrubla, J., Warbrick, T., Butz, M., Neuner, I., Shah, N.: Time-frequency analysis of resting state and evoked EEG data recorded at higher magnetic fields up to 9.4 T. The Journal of Neuroscience Methods 255, 1–11 (2015) 10.1016/j.jneumeth.2015.07.011

Abbasi, O., Hirschmann, J., Schmitz, G., Schnitzler, A., Butz, M.: Rejecting deep brain stimulation artefacts from MEG data using ICA and mutual information. The Journal of Neuroscience Methods 268, 131–141 (2016) 10.1016/j.jneumeth.2016.04.010

Alonso, J.F., Poza, J., Ma nanas, M.A., Romero, S., Fernández, A., Hornero, R.: Meg connectivity analysis in patients with Alzheimer’s disease using cross mutual information and spectral coherence. Annals of Biomedical Engineering 39(1), 524–536 (2011) 10.1007/s10439-010-0155-7

Arieli, A., Sterkin, A., Grinvald, A., Aertsen, A.: Dynamics of ongoing activity: Explanation of the large variability in evoked cortical responses. Science 273(5283), 1868–1871 (1996) 10.1126/science.273.5283.1868

Ala-Salomäki, H., Kujala, J., Liljeström, M., Salmelin, R.: Picture naming yields highly consistent cortical activation patterns: Test–retest reliability of magnetoencephalography recordings 227, 117651 (2021) 10.1016/j.neuroimage.2020.117651

Bonita, J.D., Ambolode, L.C.C., Rosenberg, B.M., Celluci, C.J., Watnabe, T.A.A., Rapp, P.E., Albano, A.M.: Time domain measures of inter-channel EEG correlations: a comparison of linear, nonparametric and nonlinear measures. Cognitive Neurodynamics 8, 1–15 (2014) 10.1007/s11571-013-9267-8

Belghazi, M.I., Baratin, A., Rajeshwar, S., Ozair, S., Bengio, Y., Courville, A., Hjelm, D.: Mutual information neural estimation. Proceedings of Machine Learning Research 80, 531–540 (2018)

Cellucci, C.J., Albano, A.M., Rapp, P.E.: Statistical validation of mutual information calculations: Comparison of alternative numerical algorithms. Physical Review E 71(6), 066208 (2005) 10.1103/PhysRevE.71.066208

Czyz, P., Grabowski, F., Vogt, J., Beerenwinkel, N., Marx, A.: Beyond normal: On the evaluation of mutual information estimators. Advances in Neural Information Processing Systems 36 (2024)

Cohen, M.X.: Analyzing Neural Time Series Data: Theory and Practise, 1st edn. MIT Press, Boston, Massachusetts, USA (2014). 10.7551/mitpress/9609.001.0001

Chen, F., Xu, J., Gu, F., Yu, X., Meng, X., Qiu, Z.: Dynamic process of information transmission complexity in human brains. Biological Cybernetics 83(4), 355–366 (2000) 10.1007/s004220000158

Doane, D.: Aesthetic frequency classifications. The American Statistician 30(4) (1976) 10.1080/00031305.1976.10479172

Darbellay, G.A., Vajda, I.: Estimation of the information by an adaptive partitioning of the observation space. IEEE Transactions on Information Theory 45(4), 1315–1321 (1999) 10.1109/18.761290

Fraser, A.M., Swinney, H.L.: Independent coordinates for strange attractors from mutual information. Physical Review A 33(2), 1134–1140 (1986) 10.1103/PhysRevA.33.1134

Gao, S., Ver Steeg, G., Galstyan, A.: Estimation of mutual information for strongly dependent variables. Proceedings of Machine Learning Research 38, 277–286 (2015)

Hoffmann, S., Falkenstein, M.: The correction of eye blink artefacts in the EEG: A comparison of two prominent methods. PLOS ONE 3(8) (2008) 10.1371/journal.pone.0003004

Ince, R.A.A., Giordano, B.L., Kayser, C., Rousselet, G.A., Gross, J., Schyns, P.G.: A statistical framework for neuroimaging data analysis based on mutual information estimated via a Gaussian copula. Human Brain Mapping 38(3), 1541–1573 (2017) 10.1002/hbm.23471

Ince, R.A.A., Rijsbergen, N.J., Thut, G., Rousselet, G.A., Gross, J., Panzeri, S., Schyns, P.G.: Tracing the flow of perceptual features in an algorithmic brain network. Scientific Reports 5(17681) (2015) 10.1038/srep17681

Jeong, J., Gore, J.C., Peterson, B.S.: Mutual information analysis of the EEG in patients with alzheimer’s disease. Clinical Neurophysiology 112(5), 827–835 (2001) 10.1016/s1388-2457(01)00513-2

Khan, S., Bandyopadhyay, S., Ganguly, A.R., Saigal, S., Erickson, D.J., Protopopescu, V., Ostrouchov, G.: Relative performance of mutual information estimation methods for quantifying the dependence among short and noisy data. Physical Review E 76(2) (2007) 10.1103/PhysRevE.76.026209

Kolmogorov, A.: On the Shannon theory of information transmission in the case of continuous signals. IRE Transactions on Information Theory 2(4), 102–108 (1956) 10.1109/TIT.1956.1056823

Kraskov, A., Stögbauer, H., Grassberger, P.: Estimating mutual information. Physical Review E 69(6), 066138 (2004) 10.1103/PhysRevE.69.066138

Laarne, P., Amnell, E., Zaidan, M.A., Mikkonen, S., Nieminen, T.: Exploring non-linear dependencies in atmospheric data with mutual information. Atmosphere 13(7) (2022) 10.3390/atmos13071046

Linfoot, E.H.: An informational measure of correlation. Information and Control 1(1), 85–89 (1957) 10.1016/S0019-9958(57)90116-X

Laarne, P., Zaidan, M.A., Tuomo, N.: ennemi: Non-linear correlation detection with mutual information. SoftwareX 14 (2021) 10.1016/j.softx.2021.100686

Moon, Y.-I., Rajagopalan, B., Lall, U.: Estimation of mutual information using kernel density estimators. Physical Review E 52, 2318–2321 (1995) 10.1103/PhysRevE.52.2318

Nagel, D., Diez, G., Stock, G.: Accurate estimation of the normalized mutual information of multidimensional data. The Journal of Chemical Physics 161(5) (2024) 10.1063/5.0217960

Na, S.H., Jin, S.-H., Kim, S.Y., Ham, B.-J.: EEG in schizophrenic patients: mutual information analysis. Clinical Neurophysiology 113(12), 1954–1960 (2002) 10.1016/s1388-2457(02)00197-9

Ostwald, D., Porcaro, C., Bagshaw, A.: An information theoretic approach to EEG–fMRI integration of visually evoked responses. NeuroImage 49(13), 498–516 (2010) 10.1016/j.neuroimage.2009.07.038

Ostwald, D., Porcaro, C., Bagshaw, A.: Information theoric approaches to functional neuroimaging. Magnetic Resonance Imaging 29(10), 1417–1428 (2011) 10.1016/j.mri.2011.07.013

Ostwald, D., Porcaro, C., Bagshaw, A.: Voxel-wise information theoric EEG-fMRI feature integration. NeuroImage 55(3), 1270–1286 (2011) 10.1016/j.neuroimage.2010.12.029

Papana, A., Kugiumtzis, D.: Evaluation of mutual information estimators for time series. International Journal of Bifurcation and Chaos 19(12), 4197–4215 (2009) 10.1142/S0218127409025298

Panzeri, S., Magri, C., Logothetis, N.K.: On the use of information theory for the analysis of the relationship between neural and imaging signals. Magnetic Resonance Imaging 26(7) (2008) 10.1016/j.mri.2008.02.019

Rousselet, G.A., Ince, R.A.A., Rijsbergen, N.J., Schyns, P.G.: Eye coding mechanisms in early human face event-related potentials. Journal of Vision 14(13) (2014) 10.1167/14.13.7

Ross, B.C.: Mutual information between discrete and continuous data sets. PLOS ONE 9(2) (2014) 10.1371/journal.pone.0087357

Shannon, C.E.: A mathematical theory of communication. The Bell System Technical Journal 27(3), 379–423 (1948) 10.1002/j.1538-7305.1948.tb01338.x

Silverman, B.W.: Density Estimation for Statistics and Data Analysis, 1st edn. Routledge, London, UK (1998). 10.1002/bimj.4710300745

Steuer, R., Kurths, J., Daub, C.O., Weise, J., Selbig, J.: The mutual information: Detecting and evaluating dependencies between variables. Bioinformatics 18(2) (2002) 10.1093/bioinformatics/18.suppl_2.S231

Scott, D.W., Terrell, G.R.: Biased and unbiased cross-validation in density estimation. Journal of the American Statistical Associatio 82(400), 1131–1146 (1987)

Sun, C., Yang, F., Wang, C., Wang, Z., Zhang, Y., Ming, D., Du, J.: Mutual information-based brain network analysis in post-stroke patients with different levels of depression. Frontiers in Human Neuroscience 12, 285 (2018) 10.3389/fnhum.2018.00285

Tuomaala, S., Autti, S., Cotroneo, S.F., Lioumis, P., Renvall, H., Liljestrom, M.: Automated speech artefact removal from MEG data utilizing facial gestures and mutual information. Imaging Neuroscience 3 (2025) 10.1162/imag_a_00545

Thuraisingham, R.A.: Estimating electroencephalograph network parameters using mutual information. Brain Connectivity 8(5), 311–317 (2018) 10.1089/brain.2017.0529

Virtanen, P., Gommers, R., Oliphant, T.E., Haberland, M., Reddy, T., Cournapeau, D., Burovski, E., Peterson, P., Weckesser, W., Bright, J., van der Walt, S.J., Brett, M., Wilson, J., Millman, K.J., Mayorov, N., Nelson, A.R.J., Jones, E., Kern, R., Larson, E., Carey, C.J., Polat, İ., Feng, Y., Moore, E.W., VanderPlas, J., Laxalde, D., Perktold, J., Cimrman, R., Henriksen, I., Quintero, E.A., Harris, C.R., Archibald, A.M., Ribeiro, A.H., Pedregosa, F., van Mulbregt, P., SciPy 1.0 Contributors: SciPy 1.0: Fundamental Algorithms for Scientific Computing in Python. Nature Methods 17, 261–272 (2020) 10.1038/s41592-019-0686-2

Xu, J., Liu, Z., Yang, Q.: Information transmission in human cerebral cortex. Physica D: Nonlinear Phenomena 106(3–4), 363–374 (1997) 10.1016/S0167-2789(97)00042-0

Yin, Z., Li, J., Zhang, Y., Ren, A., Von Meneen, K.M., Huang, L.: Functional brain network analysis of schizophrenic patients with positive and negative syndrome based on mutual information of EEG time series. Biomedical Signal Processing and Control 31, 331–338 (2017) 10.1016/j.bspc.2016.08.013

